# Thermal tolerance and survival are modulated by a natural gradient of infection in differentially acclimated hosts

**DOI:** 10.1101/2023.12.15.571921

**Authors:** Jérémy De Bonville, Ariane Côté, Sandra A. Binning

## Abstract

Wild ectotherms are exposed to multiple stressors, including parasites, which can affect their responses to environmental change. Simultaneously, unprecedented warm temperatures are being recorded worldwide, increasing both the average and maximum temperatures experienced in nature. Understanding how ectotherms, such as fishes, will react to the combined stress of parasites and higher average temperatures can help predict the impact of extreme events such as heat waves on populations. The critical thermal method (CTM), which assesses upper (CT_max_) and lower (CT_min_) thermal tolerance, is often used in acclimated ectotherms to help predict their tolerance to various temperature scenarios. Yet, few studies have characterized the response of naturally infected fish to extreme temperature events or how acute thermal stress affects subsequent survival. We acclimated naturally infected pumpkinseed sunfish (*Lepomis gibbosus*), to four ecologically relevant temperatures (10, 15, 20 and 25°C) and one future warming scenario (30°C) for three weeks, before measuring CT_max_ and CT_min_. We also assessed individual survival the week following CT_max_. Interestingly, trematode parasites causing black spot disease were negatively related to CT_max_, suggesting that heavily infected fish are less tolerant to acute warming. Moreover, fish infected with yellow grub parasites showed decreased survival in the days following CT_max_ implying that the infection load has negative survival consequences on sunfish during extreme warming events. Our findings indicate that parasite infection and high prolonged average temperatures can affect fish thermal tolerance and survival, emphasizing the need to better understand the concomitant effects of stressors on health outcomes in wild populations.

**Summary statement:** This study shows that parasites influence thermal tolerance and survival of fish, suggesting that such stressors are important to increase the ecological relevance of thermal tolerance studies of wild animals.

## INTRODUCTION

Wildlife is experiencing increases in average daily temperatures, wider daily and seasonal thermal variations and an increase in the frequency, duration and intensity of extreme weather events, such as heat waves and cold snaps (Meehl & Tebaldi, 2004; Seneviratne et al., 2014; IPCC, 2022). In ectotherms, temperature directly affects the rate of physiological functions, such as enzyme activity and energy metabolism (Fry, 1971; Clark, 2017). Thus, acute changes to the thermal environment are expected to have detrimental consequences for wild ectotherms, including increases in mortality, elevated stress levels and altered immune functions (Roth et al., 2010; Genin et al., 2020; Jørgensen et al., 2022). Although some aquatic animals are regularly exposed to rapid temperature changes, for instance, while crossing the thermocline during vertical migrations or due to diurnal temperature fluctuations in shallow waters or intertidal zones (Bates & Morley, 2020; Desforges et al., 2023), exposure of shallow aquatic species to rapid thermal changes will likely be intensified due to increases in average surface water temperatures (IPCC, 2022). Thus, estimating the susceptibility of ectotherms to acute warming events is crucial to understand the risk posed by global climate change to wild populations. Lower thermal tolerance is also relevant physiologically and ecologically. Rapid cooling events occur naturally, due to cold snaps and marine upwelling events (Donaldson et al., 2008; Lehmann & Myrberg, 2008), or anthropogenically, due to industrial runoff (Michie et al., 2020), possibly leading to mass mortality events (Reid et al., 2022). With climate change causing more thermal variability, sudden cold snaps could become more widespread in the wild (ELPC, 2019), highlighting the need to consider lower thermal tolerance when studying how climate change will affect wild fish populations (Reid et al., 2022). Many experimental studies have attempted to determine species’ thermal limits and acclimation to rapidly changing temperatures. However, the focus on temperature as the sole stressor ignores the reality of climate change, whereby multiple stressors are expected to impact wild populations simultaneously (Pirotta et al., 2022).

Indeed, wild animals experience multiple biotic and abiotic stressors concurrently, which may interact. For instance, hypoxia generally exerts antagonistic effects on thermal tolerance (Andreassen et al., 2022), possibly due to limitation of oxygen supply to tissues at warmer temperatures, limiting aerobic scope (Ern et al., 2016). However, very few studies have explored the ways that biotic stressors such as parasite infection, interact with heat stress. Despite being ubiquitous in most natural environments, parasites are seldom considered in studies of wild animal performance (Chrétien et al., 2022). This is a potentially serious oversight since ignoring infection can lead to erroneous conclusions at individual host, population and community levels (Timi & Poulin, 2020). Furthermore, some parasites, such as helminths, are expected to thrive with increasing temperatures caused by climate change : warmer water temperatures can enhance parasite growth (Macnab & Barber, 2012) and virulence (Thomas & Blanford, 2003) making host fish populations particularly at risk (Marcogliese, 2008) (but see Wood et al., 2023). Parasites can impose a range of physiological costs on hosts including increasing host metabolic demands (Thambithurai et al., 2022). Furthermore, thermoreceptors in the skin allow fish to detect and react to changing temperatures through endocrine signalling (Haesemeyer, 2020). Damage to the skin caused by parasites could potentially affect how fish react to changes in temperature. Thus, infection could impair an individual’s ability to withstand acute thermal stress by constraining oxygen uptake or impairing sensorial functions. Indeed, experimental infection with Chytrid fungus (*Batrachochytrium dendrobatidis*), led to a 4°C decrease in the thermal tolerance of frogs (*Litoria spenceri*) (Greenspan et al., 2017). Similarly, upper thermal tolerance has been negatively related to parasite load in naturally infected juvenile brown trouts (*Salmo trutta*) (Bruneaux et al., 2017), bluegills (*Lepomis macrochirus*) and longear sunfishes (*Lepomis megalotis*) (Lutterschmidt et al., 2007). Interestingly, the impact of infection on thermal tolerance appears to be parasite species-specific. In three cyprinid species, a higher intensity of trematode metacercaria (e.g. black spot disease) did not lead to a change in upper or lower thermal tolerance (Hockett & Mundahl, 1989). Likewise, Powell & Gamperl (2016) did not find differences in the upper thermal tolerance of Atlantic cod (*Gadus morhua*) infected with microsporidia (*Loma morhua*) compared to uninfected conspecifics. These mixed results may be related to the specific physiological costs of parasites on hosts. Unfortunately, the few studies that have combined measures of parasite infection and upper thermal tolerance focus on infection with a single parasite species at a given temperature, without evaluating how these relationships are affected by different acclimation temperatures. Shedding light on the relationship between parasites and host thermal tolerance in the context of climate change is essential for improving our understanding of ecological consequences infection could exert on the host, from the individual to the community level.

The critical thermal method (CTM) can be used to assess upper (CT_max_) and lower (CT_min_) thermal tolerance of animals acclimated at a given temperature (Beitinger et al., 2000; Morgan et al., 2018). Knowing the upper and lower physiological thermal thresholds of a species can help us predict which are more at risk of mass mortality events during heat waves or cold snaps (Genin et al., 2020; Reid et al., 2022; Desforges et al., 2023). Although CTM is meant to be a sublethal measure, researchers tend to assess survival only up to 24 hours after the test (Cowan et al., 2023; Stewart et al., 2023). Yet, in nature, the negative effects of thermal stress can persist for days following the onset of a heating (Suryan et al., 2021) or a cooling (Reid et al., 2022) event. Indeed, extreme events exacerbated by global change are likely to pose a greater threat to the survival of wild populations compared to average temperature increases (Vasseur et al., 2014). Thus, assessing thermal tolerance and individual survival in the days following a thermal challenge can help shed light on a species risk to climate change and improve conservation goals for aquatic ectotherms.

Here, we assessed the upper and lower thermal tolerance of a naturally infected freshwater fish (pumpkinseed sunfish; *Lepomis gibbosus* (Linnaeus 1758)) acclimated to ecologically relevant temperatures and documented their survival for one week following an acute thermal maximum trial (CT_max_). By working with wild caught fish housed in a flow-through system and considering their natural gradient of infection to assess their thermal tolerance in a lab controlled environment, we hope to increase ecological relevance of results obtained in a laboratory (Turko et al., 2023). We expected acclimation temperature to affect both CT_max_ and CT_min_ and that parasite intensity would also be related to thermal tolerance. We also expected survival to be related to infection intensity and acclimation temperature. We predicted that fish with a higher parasite load would have lower thermal tolerance (lower CT_max_ and higher CT_min_) and that heavily infected individuals would have lower survival rates. We further predicted that the effects of parasites would be more apparent at warmer temperatures, as these concomitant stressors could interact to negatively affect outcomes (Teffer et al., 2019). To our knowledge, this is the first study to address how temperature acclimation and parasite infection interact while evaluating thermal tolerance and survival following acute thermal stress in ectotherms.

## MATERIALS AND METHODS

### Model species

Pumpkinseed sunfish (Family: Centrarchidae) are native to temperate and boreal lakes of eastern North America and mostly inhabit shallow vegetated littoral zones (Scott & Crossman, 1974). Summer temperatures in their native range can vary from 22 to 31°C, depending on the lake characteristics and region (Tomeček et al., 2007). The CT_max_ of pumpkinseeds has been estimated around 35°C for fish acclimated at 20°C and up to 39°C for those acclimated at 30°C (Becker & Genoway, 1979; Rooke et al., 2017). While the CT_min_ of pumpkinseeds is comparatively understudied, Becker et al. (1977) found that fish acclimated to 15°C had an average CT_min_ of 1.7°C.

In the Laurentian region of Quebec where this study took place, *L. gibbosus* are abundant and frequently infected by trematodes causing blackspot disease (*Uvulifer sp., Apophallus sp.)*, bass tapeworms (Cestoda: *Proteocephalus ambloplitis*), as well as yellow grubs (Trematoda: *Clinostomum marginatum*) (Scott & Crossman, 1974; Guitard et al., 2022) (Fig. 1). Previous work in these populations has demonstrated significant relationships between metabolic rates and parasite infection at the cellular (Mélançon et al., 2023) and whole organism levels (Guitard et al., 2022; Thambithurai et al., 2022). The wide thermal tolerance of this species as well as high prevalence and intensities of infection make it an ideal candidate to evaluate how both factors interact in wild populations.

**Fig. 1.**
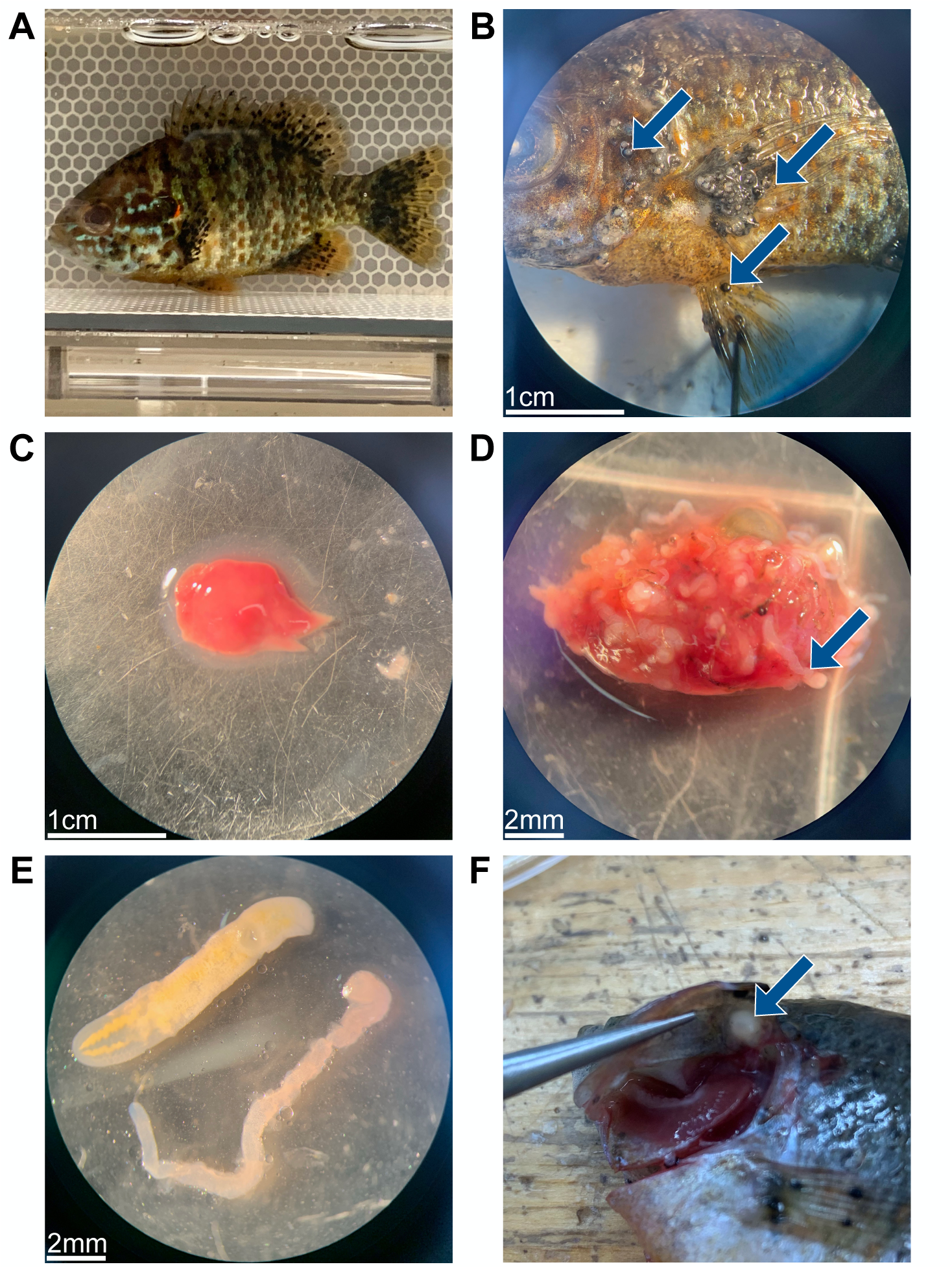
Photographs showing parasites of their host (*L. gibbosus*). (A) Black spot trematodes (*Uvulifer sp., Apophallus sp.*) on a live sunfish. (B) Blackspots shown on the body, pectoral and pelvic fins. (C) A fish liver with no parasites. (D) A fish liver infected by bass tapeworms (*P. ambloplitis*). (E) comparison between a yellow grub (*C*. *marginatum*) on the top left and a bass tapeworm on the bottom right. (F) Encysted yellow grub on the opercular flap of a pumpkinseed sunfish (F).

### Fish collection and husbandry

We collected fish in 2020 and 2021 in Lake Cromwell (SBL, QC, Canada; 45.98898°N, −74.00013°W) located on the Université de Montréal’s Station de biologie des Laurentides (SBL). Baited minnow traps were set at different sites along the littoral zone of the lake for 1-2 hours after which selected fish were collected and brought back to the lab within 30 minutes (400m walk from lake to lab). Fish underwent a hydrogen peroxide dip (2.5 ml of 3% H_2_O_2_ per litre of freshwater) for 30 minutes to limit the proliferation of external pathogens. Fish were then distributed among three flow-through 600L (215×60×60 cm, L×W×H) Living Stream tanks (Frigid Units Inc.) separated into three sections and provided with artificial plants and PVC tubes for shelter. Water was pumped from the nearby Lake Croche (45.99003°N, −74.00567°W; upstream of Lake Cromwell), and filtered through a sand pool filter, oxygenated in a collecting basin, and sterilized with UV light before entering the tanks. 24 hours following collection, fish were weighed, measured and individually marked with coloured visible implant elastomer tags (VIE, NorthWest Marine Technology Inc.) on either side of the dorsal fin with a 29-gauge needle. Fish were fed to satiety twice a day with frozen bloodworms (Hikari^®^) and tanks were siphoned daily.

In 2020, we collected 102 fish in September (mean ± s.d.: total length (TL) = 7.3±0.9 cm, mass = 6.85±2.51 g). In 2021, we collected 113 fish in June for the summer group (mean ± s.d.: TL = 7.7±0.8 cm, mass = 7.83±2.60 g) and 100 in September for the fall group (mean ± s.d.: TL= 8.1±0.9 cm, mass = 9.59±3.60 g). In fall 2020 and 2021, lighting followed a typical fall diel cycle (12L:12D) and a summer cycle (14L:10D) for fish collected in summer 2021.

Fish collection, treatment and experiments were conducted with approval from the Université de Montréal’s animal care committee (Comité de déontologie de l’expérimentation sur les animaux; certificate numbers 20-042 & 21-028) and the Ministère des forêts, de la faune et des parcs (permit numbers 2020-07-10-1730-15-S-P and 2021-05-18-1833-15-S-P).

### Acclimation treatments

We acclimated fish at 4 temperatures reflecting the natural conditions found in Lake Cromwell (10, 15, 20 and 25°C) and to one temperature representing a future climate change scenario (30°C). Historic records and colleagues (Roxane Maranger, Alice Parkes) working on Lake Cromwell were consulted in 2020 to help establish accurate acclimation temperatures. We also deployed a temperature logger (HOBO Pendant^®^ MX2201, USA, accuracy : ± 0.5 °C, resolution : 0.04 °C) 1m below the surface from June 2021-August 2022 to measure seasonal and daily variations in the littoral zone of Lake Cromwell to confirm the thermal profile experienced by these fish (Fig. 2).

**Fig. 2.**
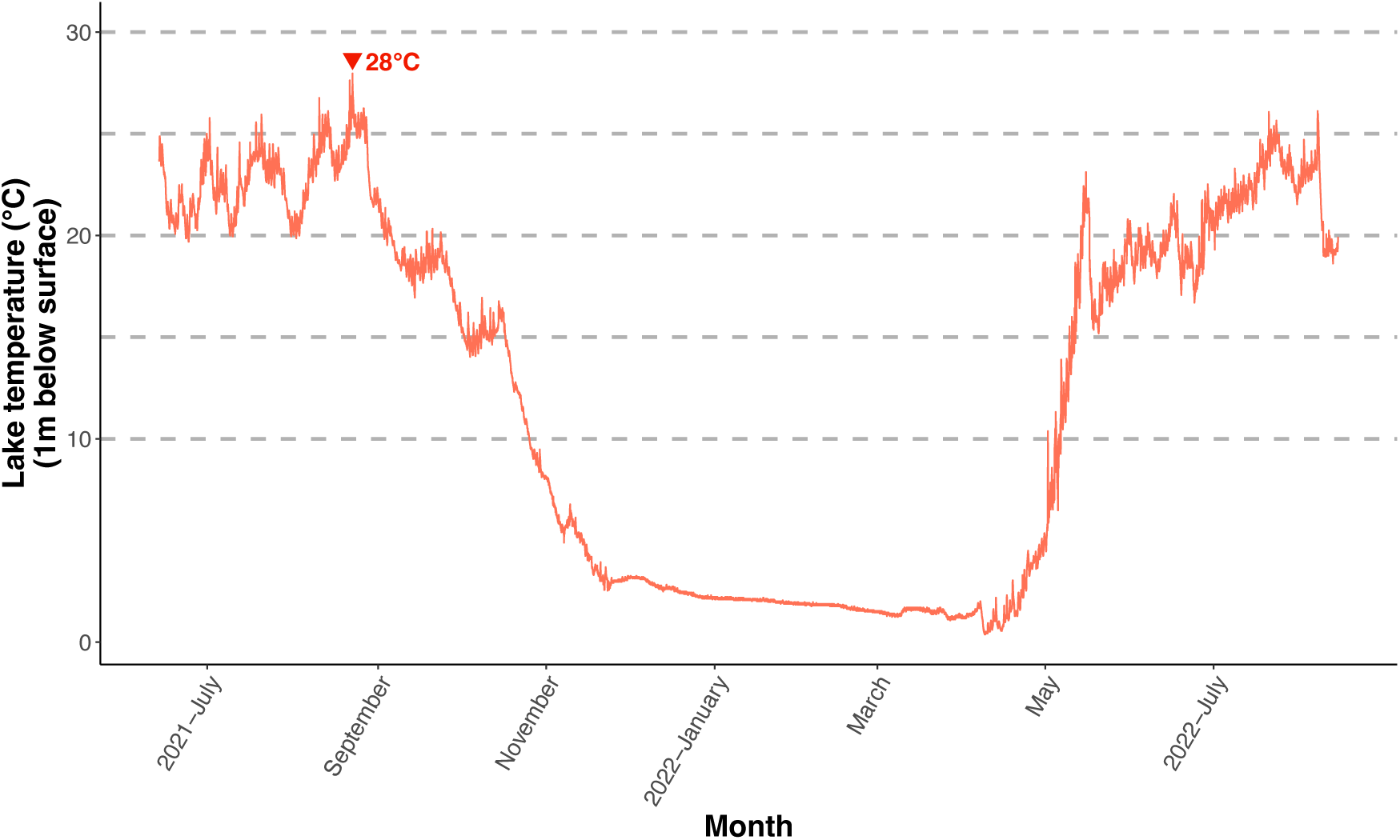
Lake Cromwell temperature readings (°C) taken with a HOBO logger (HOBO Pendant^®^ MX2201) placed 1 meter below water surface. Data was collected from 2021-06-13 to 2022-08-15 every 10 minutes. Dashed lines represent the acclimation temperatures chosen for this study. The red inverted triangle represents the warmest temperature recorded (27.97°C) on 2021-08-22.

**Fig. 3.**
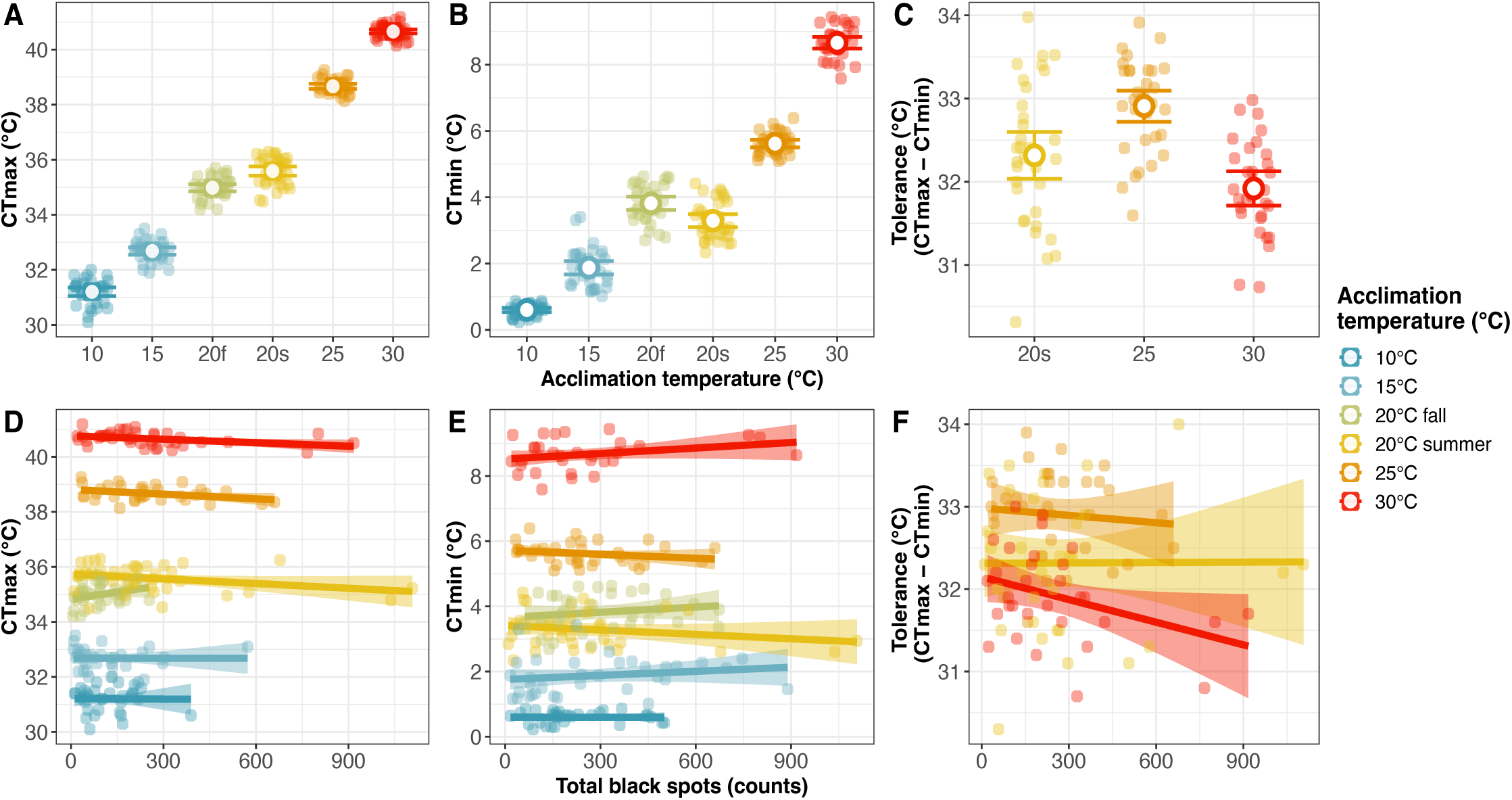
Thermal traits of pumpkinseed sunfish across acclimation temperatures and the influence of total black spots. (A) CT_max_, (B) CT_min_ and (C) thermal tolerance (CT_max_ - CT_min_) of pumpkinseed sunfish acclimated at 10, 15 and 20°C in the fall and at 20, 25 and 30°C in the summer. Each coloured circle represents an individual, larger white circles represent the means of the treatment and error bars show the 95% confidence interval. Linear regressions estimating the effect of total parasite density on (D) CT_max_, (E) CT_min_ and (F) tolerance are shown for each acclimation treatment with dots as individuals and the shaded zone as the 95% confidence interval for the regression.

For the fall groups (2020 and 2021), fish were acclimated to 10, 15 and 20°C while acclimation temperatures of 20, 25 and 30°C were chosen for summer experiments (2021). In Lake Cromwell, temperatures of 25°C represent warm summer temperatures, 20°C represents lower summer temperatures, 15°C represents early fall/late spring temperatures and 10°C represents late fall/early spring (Fig. 2). During a summer heat wave, lake temperatures can reach up to 28°C. Thus, the 30°C acclimation group represents a long thermal acclimation to heat-wave temperatures to which we added 2°C to simulate a future climate change scenario (IPCC, 2022). One day following tagging, water temperature in the flow-through tanks was increased or decreased by 1 – 1.5°C per day, depending on treatment, until reaching the desired acclimation temperature. Temperature in the tanks was monitored twice daily with hand-held thermometers, and logged through the duration of the acclimation period and experiments with a HOBO MX2201 at 10-minute intervals (Fig. S1). Temperature was maintained by controllers (Inkbird ITC-308S, China) activating either a cooling (EK20 immersion cooler, Thermo ScientificTM, USA) or a heating immersion 1500W or 2000W coil (GESAIL, China) to ensure the tanks remained at the desired acclimation temperature (± 0.2°C). Depending on the temperature, water inflow in the tanks ranged from 4.8 L/h to 22.8 L/h. This ensured an inflow of oxygenated water and a complete water change between 125h to 26h, maintaining a stable temperature and high water quality high throughout acclimation. Natural temperature changes in Lake Croche, where the tank water is pumped from, or power outages caused spikes in temperature during the acclimation periods (Fig. S2A: Oct-11, C: Sep-23). Pumps (ECO-396, EcoPlus®, China) were used to circulate the water to ensure homogenous mixing and air bubblers were deployed throughout the tank to maintain oxygen saturation.

Fish were acclimated for three weeks to allow thermal compensation before being tested for CT_max_ (Becker & Genoway, 1979; Dent & Lutterschmidt, 2003). A minimum of a 3-week acclimation period was also necessary before conducting trials, as it takes approximately 21 days for trematode cercaria to encyst and be visible as black spots on the fish (Berra & Au, 1978).

### CT_max_ & CT_min_ trials

CT_max_ and CT_min_ trials were conducted in a test arena (93 x 42 x 44 cm) containing 50L of water connected to a heating/cooling sump (50 x 30 x 26 cm) which contained 25L, for a combined volume of 75L (Fig. S2). In each corner, there were air bubblers and an inflow or outflow of water to ensure continuous mixing and homogenous temperatures between both tanks.

Fish were fed in the morning the day before the trial, then fasted for at least 24 hours before the start. Fish were habituated to the arena overnight at the same temperature as their acclimation tank, with temperature maintained stable by controllers (Inkbird ITC-308S, China). Trials were conducted the next day, starting between 09:12 and 14:47. Two to three trials were done for each acclimation group (number of fish / trial = 8-18).

Our CTM protocol followed methods described in Morgan et al. (2018). To begin the trial, two 1000W immersion coil heaters (GESAIL, China) were turned on in the sump for CT_max_. For CT_min_, a cooling coil was used (EK20 immersion cooler, Thermo Scientific^TM^, USA) as well as 1L ice blocks. Temperature was increased or decreased following a desired rate of 0.3°C min^-1^ (mean ± s.d.: CT_max_ = 0.26 ± 0.04 °C min^-1^, CT_min_ = −0.3 ± 0.1 °C min^-1^) (view Fig. S3 for ramping curves). Similar ramping rates were used previously in CT_max_ experiments on pumpkinseeds (Becker & Genoway, 1979; Dent & Lutterschmidt, 2003). Temperature was logged throughout the trials (2020 logger : HOBO^®^ MX2203, USA, accuracy : ± 0.2 °C, resolution : 0.01 °C, 2021 logger : HOBO^®^ MX2303, accuracy : ± 0.2 °C, resolution : 0.04 °C). Following methods described in Morgan et al. (2018), we conducted preliminary trials with 3 fish collected in 2020 in Lake Cromwell to ensure changes in fish internal temperature matched the water. Fish were anesthetized with eugenol (1:10 mix, at 0.4 ml/L), had a thermocouple inserted in the deep dorsal muscle (Type K, RS PRO) and were exposed to a ramping rate of ∼0.3°C min^-1^, which confirmed that internal ramping rates followed ramping rates in the water (Fig. S4).

We defined loss of equilibrium (LOE) as the point when a fish could no longer maintain a stable upright position in the water column for 3 continuous seconds (Lutterschmidt & Hutchison, 1997; Morgan et al., 2018). LOE for CT_min_ for fish acclimated to 10 and 15°C was slightly different. As these fish reached low temperatures and became inactive, we gently poked them with a hand net. If fish did not respond to poking for 3 seconds, we considered the fish to have reached its CT_min_ (Ford & Beitinger, 2005). To obtain the CT_max_ or CT_min_ the temperature measured by the logger at the time at which the fish was taken out was used. There was a delay in the temperature measures recorded by the logger used in 2020 (response time : 7 minutes), as this model was encased in a thick plastic case and was less responsive to real-time water temperature changes. Thus, for these fish (n=102), values recorded from a digital thermometer (HI98509, HANNA Instruments, USA, accuracy : ± 0.2°C, resolution : 0.1°C) placed in the arena with the fish during the experiments were used instead.

After reaching LOE, fish were transferred to a water-filled plastic bag placed in a cooler containing water at the same temperature of the acclimation group being tested and recovered for at least 1 hour (or until they swam normally). Before being transferred back to their holding tanks, fish were identified, weighed and measured. Following trials, fish were monitored for mortality over the following week, twice a day when fish were fed, and were considered deceased when found at the bottom of the tank with an absence of opercular movements. Then, fish either underwent a CT_min_ trial (summer 2021 group acclimated to 20, 25 and 30°C) or were sacrificed in an overdose of eugenol solution for the fall 2020 and 2021 groups (1:10 mix, at 4ml/L) and frozen at −18°C for dissection to assess parasite load. Summer 2021 fish were sacrificed and preserved in the same way following CT_min_. Tolerance was measured as the difference between CT_max_ and CT_min_, for fish which went through both trials (summer 2021).

### Parasite screening

Frozen fish were thawed to assess infection load. The fish were pinned to a Styrofoam pad by the fins and observed under a stereomicroscope to count encysted black spot metacercaria on their left side (body and fins) which was doubled to obtain the total amount for each fish. A subset of *L. gibbosus* (n = 89) sampled for another project confirmed that there is no significant difference in the number of black spots present between the left and right sides of the body (non-parametric paired samples Wilcoxon test ; *p* = 0.284, Fig. S5). The amount of black spots found on the gills was added to the total amount. All fish were infected to some extent by black spots (prevalence = 100%; median = 163 parasites; range = 6 - 1107). Dissection of internal organs and muscles was conducted to look for the presence of other parasites (specifically in the gills, heart, liver, digestive tract, gonads and in the abdominal cavity). This allowed us to estimate the load of other endoparasites, which mainly consisted of bass tapeworm (*Proteocephalus ambloplitis)* found in the liver and digestive tract, as well as the yellow grub (*Clinostomum marginatum*) mostly found in the gills, heart and in the muscle. While 94.3% of fish were infected by *P. amploblites* (median = 15 parasites; range = 0-186), only 23.8% of fish were infected by *C*. *marginatum* (median = 0 parasites; range = 0 - 10) (Fig. 1 for parasite images). These prevalence values are similar to results obtained by Guitard et al. (2022) on the same population of similar size (mean ± s.d.: total length = 8.5±0.7 cm, body mass = 10.24±2.46 g) also sampled with minnow traps in July 2019 (black spot : 100%, bass tapeworm : 93%, yellow grub : 26%). Individual fish mass was corrected for the presence of parasites (as suggested by Timi & Poulin, 2020) by basing parasite weight averages on values measured by Guitard et al. (2022) on the same species.

### Statistical analyses

All data were analysed in R v.4.2.2 (http://www.R-project.org/). We used general linear models (lm function in R) to model the effect of acclimation treatment and parasite count on thermal tolerance traits (CT_max_, CT_min_ and overall tolerance). To find the best fitted model to explain each trait, fish mass (corrected for parasites), the number of black spots, the number of bass tapeworms, the number of yellow grubs, the acclimation treatment, the interaction between mass and acclimation treatment as well as all the interactions between parasite groups and acclimation treatment were used as predictors in the three full models. Although parasite count and fish mass were correlated (cor=0.29), collinearity of variables included in the models was tested using the variance inflation factor (VIF). As the VIF was low (<2), both variables were included in the models (Legendre & Legendre, 2012).

We used Akaike’s Information Criterion (AIC) to compare candidate models for each trait and select the most parsimonious model (Burnham et al., 2011; Thambithurai et al., 2022; Stewart et al., 2023). Parasites were included in all tested models, but mass, acclimation, and the interactions were removed in a stepwise manner. We compared general linear models using ΔAIC, AIC cumulative weights, and *R*^2^ and, for the best model, the effects of each selected predictor was assessed using the Anova() function and estimates of each predictors were evaluated using the summary() function to assess the direction of the effect, if significant. We used post-hoc contrasts of least square means using the emmeans package (Lenth, 2022) to assess differences between acclimation temperatures. We visually assessed model assumptions with diagnostic plots which were met for all models.

The packages “survival” (Therneau, 2023) and “survminer” (Kassambra et al., 2017) were used for the survival analyses, following methods from (Fox & Weisberg, 2011). A proportional hazards survival regression (coxph function from the “survival” package) was used to test the effect of three different parasite species and fish mass on survival following CT_max_ tests at three acclimation treatments (20, 25 and 30°C) (Cox, 1972). This model calculates the influence of covariables on survival probability over a set period of time (7 days post CTmax trial). The response variable (hazard ratio, HR) represents the risk of death during the survival assessment period. Thus, a HR<1 indicates that the associated predictor is protective (improved survival) and an HR > 1 increases the risk (decreases survival) (Cox, 1972; Clark et al., 2003). We tested multiple candidate models, which always included parasites, to which we added acclimation temperature, fish mass, and all 2-way interactions. Starting from the most complex model, we used likelihood ratio tests (LRT) to compare models and removed non-significant interactions and predictors (Table S2) (Therneau & Grambch, 2000, Cortese et al., 2021). The effects of each predictor were assessed using the Anova() function and hazard ratios (coefficients) of each predictors were obtained using the summary() function.

Model diagnostics were assessed for all models tested, where we checked for violation of the assumption of proportional hazards, for nonlinearity in the relationship between the log hazard and the covariates used in the models and for influential data (Fox & Weisberg, 2011). The assumptions of proportional hazard were tested using the cox.zph() and visually observed by plotting the Schoenfeld residuals with the ggcoxzph() function from the survminer package and validating that all predictors where encompassed by the 95% confidence interval lines (Fig. S6). The martingale residuals were plotted against covariates to confirm the nonlinearity assumption and influential cases were negligeable when observing residuals of the model (all dfbetas <1).

## RESULTS

### Influence of acclimation and parasites on thermal traits

Across acclimation temperature treatments (10, 15, 20, 25, 30°C), thermal tolerance ranged from 31.20 °C ± 0.46 to 40.66°C ± 0.23 for CT_max_ (Fig. 2A) and 0.60°C ± 0.18 to 8.66 °C ± 0.47 for CT_min_ (Fig. 2B). Based on the average CT_max_ value of each group, CT_max_ increased on average by 0.47°C for each degree through acclimation from 10 to 30°C (9.46°C difference).

Following model selection for all thermal traits, CT_max_ was best predicted by black spots, bass tapeworms, yellow grubs, fish mass and acclimation temperature, with no interactions (CT_max_ Model 1, Table S1A, AIC = 199.15). Acclimation temperature (R^2^=0.99, *F*(5,205) = 2790.90, *p*<0.001, Table 1 and Fig. 2A) and black spot count (R^2^=0.99, *F*(1,205)=7.94, *p* = 0.005, Table 1 and Fig. 2D)) were significantly related to CT_max_. CT_max_ was higher with increasing acclimation temperature : Tukey’s post-hoc test showed significant differences between the means of all acclimation groups (*p*<0.001) (Table S1B). Conversely, black spot count was negatively related to CT_max_ (β = −0.0005, t-value =-2.82, *p* = 0.005, Table S1C). Each black spot decreases CT_max_ by 0.0005°C: i.e. a fish with 1000 black spots has an estimated CT_max_ 0.5°C lower compared to a fish with no black spots.

**Table 1.** Relationship between thermal trait and predictors as selected by the best fitting model for each trait. Significant predictors are shown in bold.

| Response | Predictors | d.f. | F-value | p-value | $R^2_{aj}$ |
| --- | --- | --- | --- | --- | --- |
| CT<sub>max</sub><br>(n=215) | **Black spots** | 1, 205 | 7.94 | **0.005** | **0.99** |
|  | Bass tapeworm | 1, 205 | 0.46 | 0.50 |  |
|  | Yellow grubs | 1, 205 | 0.19 | 0.67 |  |
|  | Fish mass | 1, 205 | 3.40 | 0.07 |  |
|  | **Treatment** | 5, 205 | 2790.90 | **<0.001** |  |
| CT<sub>min</sub><br>(n=200) | Black spots | 1, 185 | 0.43 | 0.51 | 0.97 |
|  | Bass tapeworms | 1, 185 | 1.27 | 0.26 |  |
|  | Yellow grubs | 1, 185 | 0.21 | 0.65 |  |
|  | Fish mass | 1, 185 | 0.54 | 0.46 |  |
|  | **Treatment** | 5, 185 | 1145.01 | **<0.001** |  |
|  | **Fish mass \* Treatment** | 5, 185 | 4.16 | **0.001** |  |
| Tolerance<br>(n=100) | Black spots | 1, 93 | 1.99 | 0.38 | 0.26 |
|  | Bass tapeworms | 1, 93 | 0.08 | 0.37 |  |
|  | Yellow grubs | 1, 93 | 0.01 | 0.08 |  |
|  | Fish mass | 1, 93 | 2.26 | 0.08 |  |
|  | **Treatment** | 2, 93 | 16.19 | **<0.001** |  |

See Table S1B for model results with estimates for each coefficient tested in the models.

CT_min_ decreased on average by 0.40°C for each degree from 30 to 10°C (8.06°C difference). CT_min_ was best predicted by a combination of black spots, bass tapeworms, yellow grubs, fish mass, acclimation temperature, and an interaction between fish mass and acclimation temperature (CT_min_ Model 1, Table S1A, AIC = 275.83). In this model, acclimation temperature (R^2^=0.97, *F*(5,185) = 1145.0, *p*<0.001, Table S1B and Fig. 2B) and the interaction between mass and acclimation (R^2^=0.97, *F*(5,185)=4.16, *p* = 0.001, Table 1)) were significantly related to CT_min_. Tukey’s post-hoc test further showed significant differences in CT_min_ among all acclimation temperatures (*p*≤0.001) (Table S1B). The only significant interaction between fish mass and acclimation temperature was within the 15°C group (β = 0.105, t-value =3.41, *p* = 0.0008).

The best model to explain overall thermal tolerance included black spots, bass tapeworms, yellow grubs, fish mass and acclimation temperature, with no interaction, (Tolerance Model 1, Table S1A, AIC = 210.53). Only acclimation temperature had a significant effect on predicting tolerance (R^2^=0.26, *F*(2,93) = 16.19, *p*<0.001, Table S1B and Fig. 2C) and we found significant differences between the 20°C and 25°C groups (*p* = 0.0011), the 25°C and 30°C (*p* < 0.0001) groups, but not between the 20°C and 30°C acclimated fish (*p* = 0.099) (Table S1B).

### Influence of parasites on survival between acclimation groups

Mortality was higher in the 30°C group, as only 73.8% (31/42) of fish survived the week, as opposed to 94.6% (35/37) and 100% (34/34) for the 20°C and 25°C groups respectively (Fig. 4). Although survival ratio seems lower at warmer acclimation temperatures (Fig. 4A), as shown by its Hazard ratio (HR) >1, acclimation was not significant in explaining survival in our best fitting model (coxph model: Acclimation; HR = 1.17, *p* = 0.0896, Table 2). The amount of yellow grubs, on the other hand, was significant in explaining survival through time (coxph model: Yellow grubs; HR = 2.10, *p* = 5.55e-06, Table 2), where an increase in yellow grubs leads to a decrease in survival probability following a CTmax trial (Fig. 4B), shown by the HR>1.

**Fig. 4.**
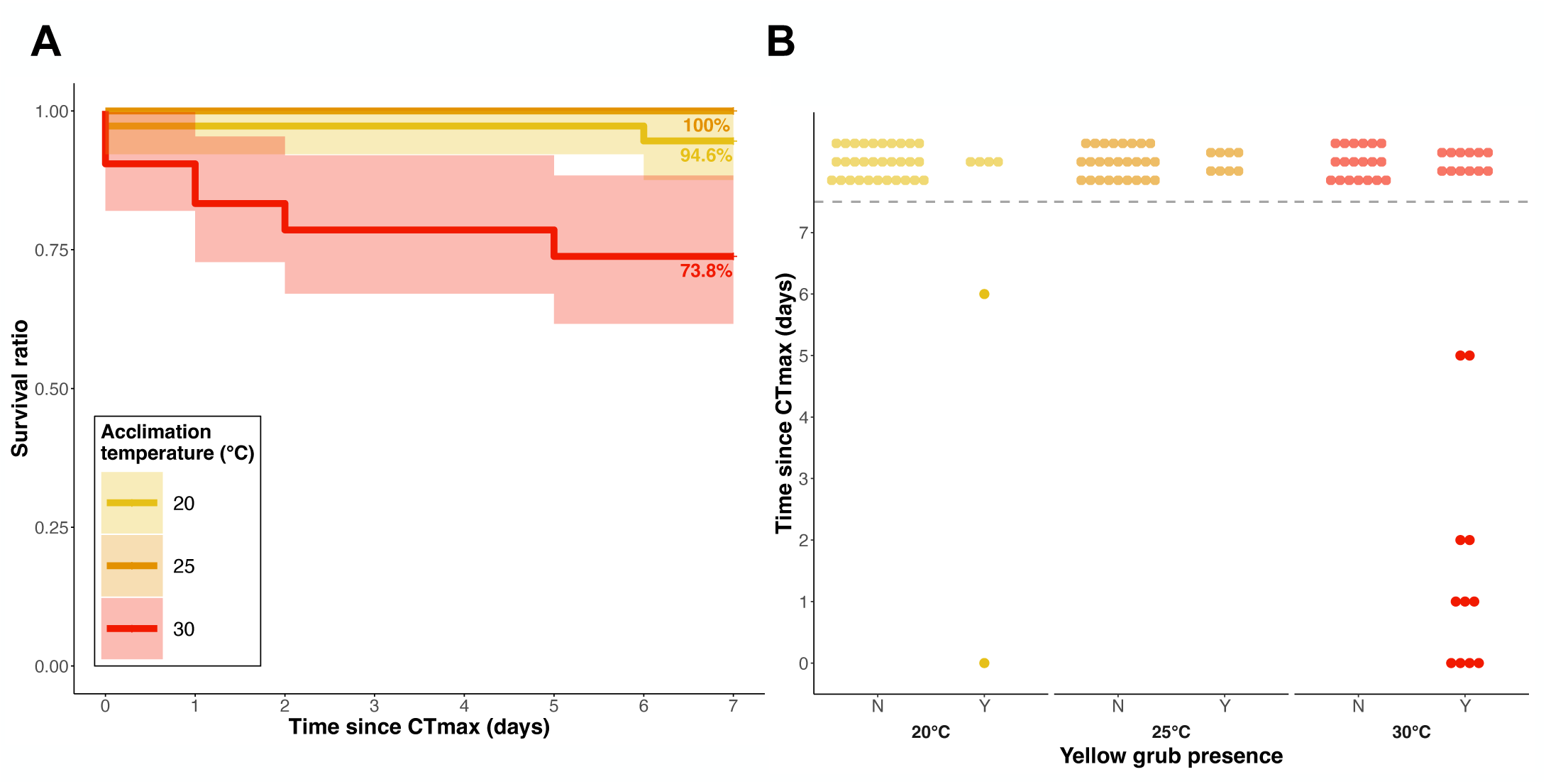
Survival of pumpkinseed sunfish in the week following a CT_max_ test across 3 temperatures (20, 25 and 30°C) and the influence of yellow grub trematodes. (A) Kaplan-Meier Kaplan-Meier survival curves for *L.gibbosus* survival post CT_max_ for 3 acclimation temperatures including the number of surviving fish per day. (B) mortality events at three acclimation temperatures according the yellow grub presence (Y) or absence (N). The shaded zone represents 95% confidence intervals in (A) and dots represent individuals in (B). Dots above the dotted line represent fish that survived the 7-day period following a CT_max_ trial. Although the model considered the parasites number as a count, they are represented here as a binomial variable for visual purposes.

**Table 2.** Results of the best fit coxph model estimating the relationships between survival and parasites, fish mass and acclimation temperature.

| Predictor | HR | 95% CI | *p*-value |
| --- | --- | --- | --- |
| Black spots | 1.00 | 1.00, 1.00 | 0.524 |
| **Yellow grubs** | **2.10** | **1.52, 2.89** | **5.55e-06** |
| Bass tapeworms | 0.97 | 0.90, 1.04 | 0.376 |
| Fish mass | 0.98 | 0.74, 1.30 | 0.892 |
| Acclimation | 1.17 | 0.98, 1.39 | 0.0896 |

In bold are the predictors with a significant effect on survival. (HR : Hazard ratio; CI : Confidence intervals)

## DISCUSSION

We investigated the relationships among thermal tolerance, parasites and acclimation to four ecologically relevant temperatures and one future climate change scenario in wild-caught pumpkinseed sunfish. We were also able to explore relationships between survival, infection and acclimation temperature following an acute thermal event. As predicted, acclimation temperature affected all thermal traits measured: CT_max_ was higher for fish acclimated at 30°C, CT_min_ was lowest when fish were at 10°C and thermal tolerance was maximized at 25°C. Interestingly, higher black spot load was related to a lower CT_max_ and survival decreased for individuals infected with yellow grubs. However, parasite load did not explain variation in CT_min_ nor tolerance.

Our results align with other thermal tolerance studies on pumpkinseeds. Under similar acclimation treatments, pumpkinseeds collected from Rice Lake and the Otonabee River (Ontario, Canada) had an average CT_max_ of approximately 30°C and 39.5°C when acclimated at 10 and 30°C respectively (Rooke et al., 2017), compared to our values of 31.2°C and 40.7°C. The small discrepancy in values is possibly explained by their slower ramping rate of 1°C every 15 minutes (0.066°C min^-1^). Fishes exposed to faster rates during trials generally have a higher CT_max_ (Illing et al., 2020; Leong et al., 2022). However, CT_max_ in zebrafish measured during slow (0.025°C min^-1^) and fast (0.3°C min^-1^) ramping rates were correlated (Åsheim et al., 2020). This slow thermal ramping resembles natural rates of water warming during heat waves experienced in Lake Cromwell. Indeed, temperatures in the littoral zone where our sunfish were collected increased from 24.66°C to 27.63°C in 2 hours (0.025°C min^-1^) during a heat wave in August (Figure S1). Thus, although we might expect lower absolute CT_max_ temperatures in fish experiencing more natural ramping rates in the wild, our laboratory results likely reflect relative differences in an individual’s upper thermal tolerance measured using fast ramping rates.

Across all acclimation temperatures, we found that CT_max_ was lower in fish that had a high intensity of black spot metacercaria. This is consistent with Lutterschmidt et al. 2007 which found that two centrarchid species (*Lepomis macrochirus* and *Lepomis megalotis*) highly infected by trematodes had lower CT_max_ than conspecifics with low parasite intensity (approximately 32°C versus 35.5°C respectively). Early stages of infection with black spot trigger an immune response in their hosts, which can be energetically demanding (Lemly & Esch, 1984), and potentially detrimental to upper thermal tolerance. The mechanisms underlying this relationship need to be further explored.

The average CT_min_ of our pumpkinseeds acclimated at 15°C (1.9°C) was similar to those in Becker et al. (1977): 1.7°C. However, we did not find any relationship between parasite load and cold tolerance. Few studies investigate CT_min_ in temperate populationsg249 cl:930, and even fewer have directly evaluated whether parasites affect CT_min_ (Hockett & Mundahl, 1989). As average temperatures are rising and winters become shorter (Wang et al., 2021), temperate fishes may become more sensitive to acute cold shocks. This can be illustrated by the stark contrast in the average CT_min_ of fish acclimated at 20°C (3.3°C), compared to those at 25°C (7.1) or 30°C (8.7), as the difference in tolerance between 25 and 30 is only 1.6°C, while it is 3.8°C between 20 and 25. In the context of climate change where average water temperatures and cold shocks are expected to increase (Reid et al., 2022), temperate fishes could be at risk to extreme events due to the rapid loss of lower thermal tolerance at warmer acclimation temperatures. Similar to CT_max_, we predicted that parasite infection would be related to increased CT_min_. Indeed, Lemly & Esch (1984) found that high intensities of black spot metacercaria decrease bluegill (*Lepomis macrochirus*) lipid content, leading to lower survival rates in overwintering fish. The fact that we found no relationship between parasite load and CT_min_ could indicate that the underlying mechanisms of CT_max_ and CT_min_ are differentially affected by parasites: infection may constrain upper thermal tolerance acutely, but limit lower thermal tolerance over longer time periods, such as during overwintering. There is a need to further explore the links between cold tolerance, overwintering survival and infection.

To our knowledge, no study has measured CT_max_ and CT_min_ on the same individuals to obtain overall thermal tolerance values for *L. gibbosus*. Here, for three summer acclimation temperatures (20, 25, 30°C) we found that tolerance was highest at 25°C and lowest at 30°C. Although CT_max_ is maximized at 30°C, individuals may be reaching a hard ceiling where acclimation to warmer temperatures will no longer increase CT_max_, possibly limiting their adaptability in a warming climate (Sandblom et al., 2016). Indeed, a simultaneous faster increase in CT_min_ resulted in a lower overall thermal tolerance for fish in this acclimation group. Interestingly, the higher thermal tolerance at 25°C suggests that this temperature provides pumpkinseed with a wide thermal window, and may be optimal for activities such as growth and reproduction. Exposure to warmer temperatures such as at 30°C may constrain aerobic scope (Jutfelt et al., 2021) or induce a stress response in individuals (Alfonso et al., 2021) as our populations do not typically reach temperatures above 28°C in the summer. Although we expected parasites to constrain tolerance across all temperatures, we found no significant relationship between infection and tolerance. This supports the idea that the underlying mechanisms constraining upper and lower thermal tolerance could be affected in different ways by parasite infection.

Although CT_max_ protocols should be non-lethal, most studies only assess survival after 1h (Ford & Beitinger, 2005), or 24h (Cowan et al., 2023; Stewart et al., 2023), or omit to mention the survival assessment period (Bennett & Beitinger, 1997; Morgan et al., 2018). Temperatures in the wild can reach or approach an individual’s CT_max_ during massive heat waves (Desforges et al., 2023) and consequences are likely felt over multiple days (Genin et al., 2020; Alfonso et al., 2021). Our results suggest that mortality can occur multiple days after a CT_max_ trial: death occurred at 1, 2, 3 and 6 days post trial for fish acclimated at 30°C and after 1 and 6 days in the 20°C group. The difference in survival rates across temperatures was best explained by the presence of yellow grubs found in fish gills, where infected fish acclimated at 30°C seemed to have lower survival rates. Although the ecology and life cycle of the yellow grub is well understood (Wang et al., 2017), research on the physiological effects of infection on hosts remains scarce. Considering these parasites target metabolically active organs, such as the heart, gills or muscles, we suggest infection with this parasite imposes an additional toll for recovery following an acute thermal challenge for individuals acclimated to temperatures that are stressful (Marcogliese, 2008). Indeed, parasites have been linked to reduced survival following heat stress in Coho salmon (*Oncorhynchus kisutch*) (Teffer et al., 2019), mosquito fish (*Gambusia affinis*) (Granath & Esch, 1983) and three-spined sticklebacks (*Gasterosteus aculeatus*) (Wegner et al., 2008). Our results suggest that summerkill events caused by heat wave events, which are expected to increase more than fourfold in the northern hemisphere, could impose considerable risk to infected fish (Meehl & Tebaldi, 2004; Seneviratne et al., 2014).

### Concluding remarks

Our study provides novel information on CT_min_ and thermal tolerance of this species, which is scarce, as well as highlights the need to consider survival when conducting thermal tolerance studies in the lab, as effects may be felt multiple days later. As certain parasites seem to have differential effects, omitting to consider them could indicate that acclimation temperature was the only determining factor explaining survival and thermal tolerance. As we are currently facing exceedingly high amounts of heatwaves, fish populations are at risk, and summer die-offs could be expected, highlighting the need to start incorporating parasite load and longer survival assessment in studies on wild fish populations.

## Acknowledgements

We would like to thank Gabriel Lanthier and the Station de Biologie staff for logistical support, and Roxane Maranger and Alice Parkes for advice and data on temperature profiles in Lake Cromwell. We also thank Rachael Morgan and Fredrik Jutfelt for advice on the experimental set-up, Charles Picard-Krashevski and Rémi Chauvette for assistance during experiments, Hyunmo Koo and Arnaud Blouin for help with dissections, and Dominique Roche, Marie Levet and Juliane Vigneault for advice on statistical analyses.

## Competing interests

No competing interests declared.

## Funding sources

This work was supported by a Natural Sciences and Engineering Research Council of Canada Discovery grant (S.A.B.) and the Canada Research Chair Program (S.A.B.). J.D.B. was supported by a doctoral grant from the Fonds de recherche du Québec - Nature et technologies.

## Data availability

Data used in this study is being prepared for archiving and will be made available in a public data repository (i.e. Figshare) upon acceptance.

## Diversity statement

We acknowledge the traditional lands of the Kanien’kehá:ka, Omàmiwinini, and Anishinabewaki First Nations on which the field and laboratory work for this project took place. During the research project, we aimed to create opportunities for students from different backgrounds when selecting undergrads as field assistants or research interns. For instance, we prioritized applications from international students and students from marginalized communities.

## Notes

### Competing Interest Statement

The authors have declared no competing interest.

